# TEExplorer: A Web Portal to Investigate TE-Epigenome Associations Across Human Cell Types

**DOI:** 10.64898/2026.02.18.706470

**Authors:** Jeffrey Hyacinthe, David R Lougheed, Guillaume Bourque

## Abstract

Approximately half of the human genome is derived from transposable elements (TEs) and several studies support the involvement of TEs in genome regulation in development, immunity and disease. We previously leveraged 4614 ChIP-seq samples from the International Human Epigenome Consortium (IHEC) EpiATLAS dataset and did a comprehensive analysis of the relationship between TEs and 6 histone marks across 57 human cell types. However, with over 6 million measurements of TE / histone mark / cell type enrichment, it was challenging to navigate the results and it was not possible to integrate them with user data. To address this, we developed a web tool, TEExplorer, which makes available TE overlaps and enrichments in an accessible and intuitive manner. The tool presents an interactive view of TE families and subfamilies, with their overlap and enrichments across histone marks and cell types. Finally, the tool allows users to upload their own ChIP-seq BED file to obtain the TE overlap and enrichment relative to random controls and compare their data with the EpiATLAS dataset. With TEExplorer, researchers with an interest in a particular TE family or subfamily, histone mark, or cell type, or those bringing their own ChIP-seq dataset, can dynamically explore and contrast hundreds of associations found within the large EpiATLAS dataset.

**Availability:** Online portal: https://teexplorer.c3g.sd4h.ca

## Introduction

Transposable elements (TEs), which cover about half of the human genome^1^, are DNA sequences with the ability to transpose and duplicate themselves. As such, they have the potential to act as regulatory units that disseminated regulatory sequences throughout our genome^2^. For instance, researchers have found that the majority of primate regulatory sequences are derived from TEs^3^, and that TEs are intertwined with the epigenome^4,5^, contain transcriptional enhancers and enhancer-like sequences^6–8^, and even contribute to infection response regulatory regions^9,10^. However, due to our limited ability to characterize TEs because of their repetitive nature^11^, much of their potential function in the genome has only recently started to be explored. Moreover, TE analysis remains limited because their importance within regulation isn’t fully understood, and their analysis requires specialised tools.

In order to spread approaches and knowledge, it is just as important to share data and results as it is to make it easily accessible and interpretable^12^. Some of the most widespread exploration tools such as GREAT^13^ and the UCSC Genome Browser^14^ or datasets such as ENCODE^15^ or GTEX^16^ leverage an online interface for ease of access, usage and sharing. The UCSC Repeat Browser^17^ is a useful tool for TE consensus sequence and interactions, however, it is limited in its ability to display interpretable associations between TEs and cell types or histone marks. The WashU Repeat Browser^18^ has more powerful display options and features, and many transcription factor Chromatin ImmunoPrecipitation Sequencing (ChIP-seq) samples, but has a limited set of histone samples and cell types.

The International Human Epigenome Consortium (IHEC)^19^ is a group of international consortiums with the overall objective to better understand the role of the human epigenome in health and disease. Recently, the consortium has made available a uniformly reprocessed dataset called EpiATLAS, which contains 4614 ChIP-seq samples from 6 histone marks across 57 cell types ^19^. In our previous study leveraging the EpiATLAS dataset^4^, we presented a comprehensive overview of the relationships between TEs, cell types, and the epigenome. This included TE overlap with histone marks and TE enrichments measured by comparing observed overlaps relative to random control overlaps. However, we could only highlight some of the most striking examples that we had observed, and we believed that there are many more noteworthy associations. We wanted to make our results available in an easily navigable and interpretable manner and promote TE consideration by making similar TE analysis easy to initiate.

Here we present TEExplorer, a web portal that makes available TE enrichment data for 57 cell types across 6 histone marks which can be plotted alongside 3 different enrichment metrics. TEExplorer facilitates users in investigating these data at the level of 60 TE families or 1,426 TE subfamilies. Notably, the portal also allows users to upload their own data for a rapid analysis of TE overlaps and enrichment, and for comparison of results to those of the large EpiATLAS dataset.

## Results

### Data and structure

To develop our web tool, we used data from EpiATLAS, which included 4614 ChIP-Seq samples from 57 cell types across 6 histone marks (activating: H3K27ac, H3K4me3, H3K4me1, H3K36me3, and repressive: H3K27me3, H3K9me3). We overlapped the peaks of all samples (and associated controls from our previous TE assessment of the EpiATLAS dataset^4^) with the coordinates of TEs and repeats (thereafter jointly referred as TEs) from RepeatMasker^20^ to obtain 6,572,151 measurements of ChIP-Seq peak overlaps with TEs. We summarized these measurements into two databases, with TEs grouped by subfamily in the first and family in the second. The TE subfamily database contained an entry for each TE subfamily per sample and its cell type, assay, observed overlap count, expected count from random simulations, total peak count in the sample and calculated metrics such as observed-expected count and fold change over expected. The TE family database was a pre-calculated aggregate of the subfamily database where we summed overlap counts within subfamilies to tally family level totals per sample and calculated the fold change metric: # *observed overlaps*⁄# *expected overlaps*. By pre-computing this database, we save computation at runtime when responding to user queries.

### An overview of TE enrichments across assays and cell types

To present all these TE enrichments, we developed a web tool with three sections: a TE Overview section for a broad look at TE families and their overall enrichments, a TE Subfamilies section for investigation of specific TE subfamilies within a selected TE family and histone mark across cell types, and an Import section allowing users to quickly obtain the TE measurements of their own data as well as to compare it with the EpiATLAS dataset (**Supplemental Fig. 1**).

The TE Overview section provides an overview of TE family enrichment across cell types and histone marks (**Fig. 1**). By default, this section displays data for all major TE families across all cell types and histone marks, but users can select specific TE families, histone marks, and cell types of interest for more specific inquiry (**Fig. 1A**). The overview first shows a bar plot of TE overlap with histone marks (**Fig. 1B**). We see for instance that H3K9me3 has the highest TE overlap (73%) while the other histone marks have between 31% and 54% overlap. By selecting only a few cell types (see **Fig 1A**), we could get the TE proportion within those cell types. For instance, the TE overlap within H3K9me3 in brain is 84% (**Fig. 1C**) compared to 71% in T cells (**Fig. 1D**). Next, we display a heatmap of the TE enrichment relative to controls (observed – expected %) for all samples matching the query parameters across the TE subfamilies within the selected families (**Fig. 1E**). By default, only the most enriched TE subfamilies are shown, but all subfamilies can optionally be displayed. We can see distinct patterns of TE subfamilies being generally enriched within some TE families depending on the assay. For instance, we observe that L1 elements are particularly enriched in H3K9me3, while L2 and MIR are more enriched for H3K27ac, H3K27me3 and H3K4me1 (**Fig. 1E**). Finally, we show a bar plot overview of the mean enrichment across TE families for each histone mark. In this example, we can see a mean enrichment of L1 in H3K9me3 (6%) and depletion in all other marks (**Fig. 1F**).

**Figure 1.**
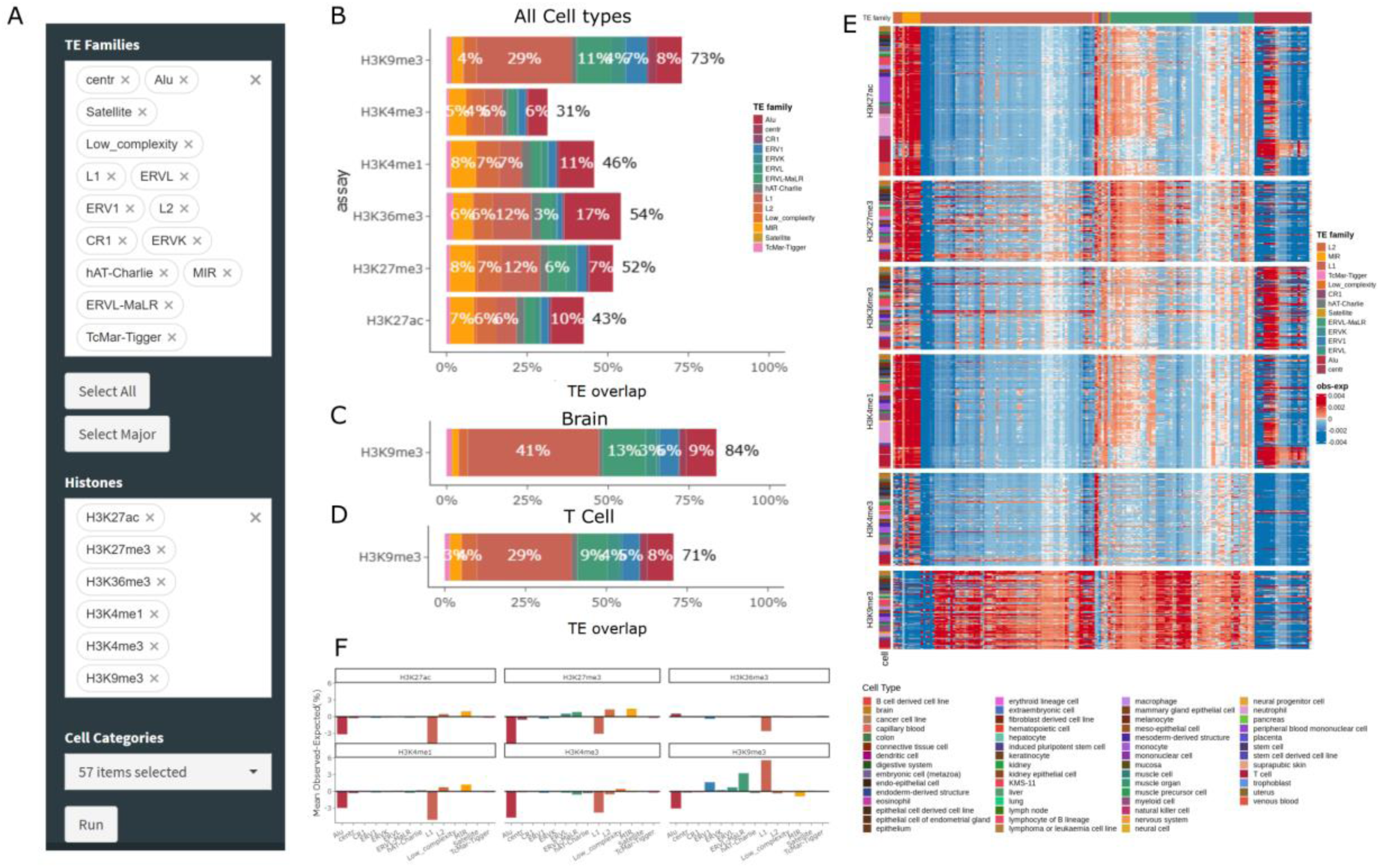
TE Overview section provides a broad overview of TE association with histone marks. **A)** Overview section input interface. TE families, assays (histone marks) and cell types to use can be chosen. **B)** Bar plot of mean TE overlap within the 6 histone marks broken down by TE family. The results shown depend on the user input query. Top shows default input of major TE families for all (57) cell types, middle for all brain only and bottom for T cells only. **C)** same as B but having only selected cell type Brain and **D)** same as B but having only selected cell type T cell. (fill color, legend in B) **E)** Heatmap of TE subfamily enrichment (Observed – Expected, obs-exp in percent) of the samples matching the user input query, left bar annotations are cell type, top annotations are TE families. **F)** Mean enrichment relative to expected (obs-exp in percent) of TE families for each selected assay.

These plots give users an idea of which TE families are most present in each cell type for the various histone marks. The heatmap enables the comparison of enrichment profiles between samples and cell types.

### Detecting the most enriched TE subfamilies within specific cell types

The TE Subfamilies section allows the investigation of a specific TE family and its subfamilies’ overlap and enrichment across cell types. The user can select a TE family and a histone mark and obtain a breakdown of the TE enrichment or observed count across cell types and subfamilies (**Fig. 2A**). This section first reports basic information of the selected TE family (instance count, subfamily count and estimated mappability, **Supplemental Fig 2A**). We also show a boxplot of that family’s enrichment or observed count for the selected histone mark across cell types. For example, after specifying the Alu family and the H3K27ac histone mark, we observe that hematopoietic cells have a median TE enrichment of 13%, the highest across cell types (**Fig. 2B**). If we look at L1 in H3K9me3, we see that the uterus, neural progenitor, and brain cell types have the highest enrichment (**Fig. 2C**). This highlights how TEs can have drastically different enrichments depending on histone mark and cell type. Note that by enabling the “include depleted TE” toggle, we allow the non-enriched TEs to be used, which can produce negative TE enrichment figures indicating depleted TEs (**Supplemental Fig 2B**).

**Figure 2.**
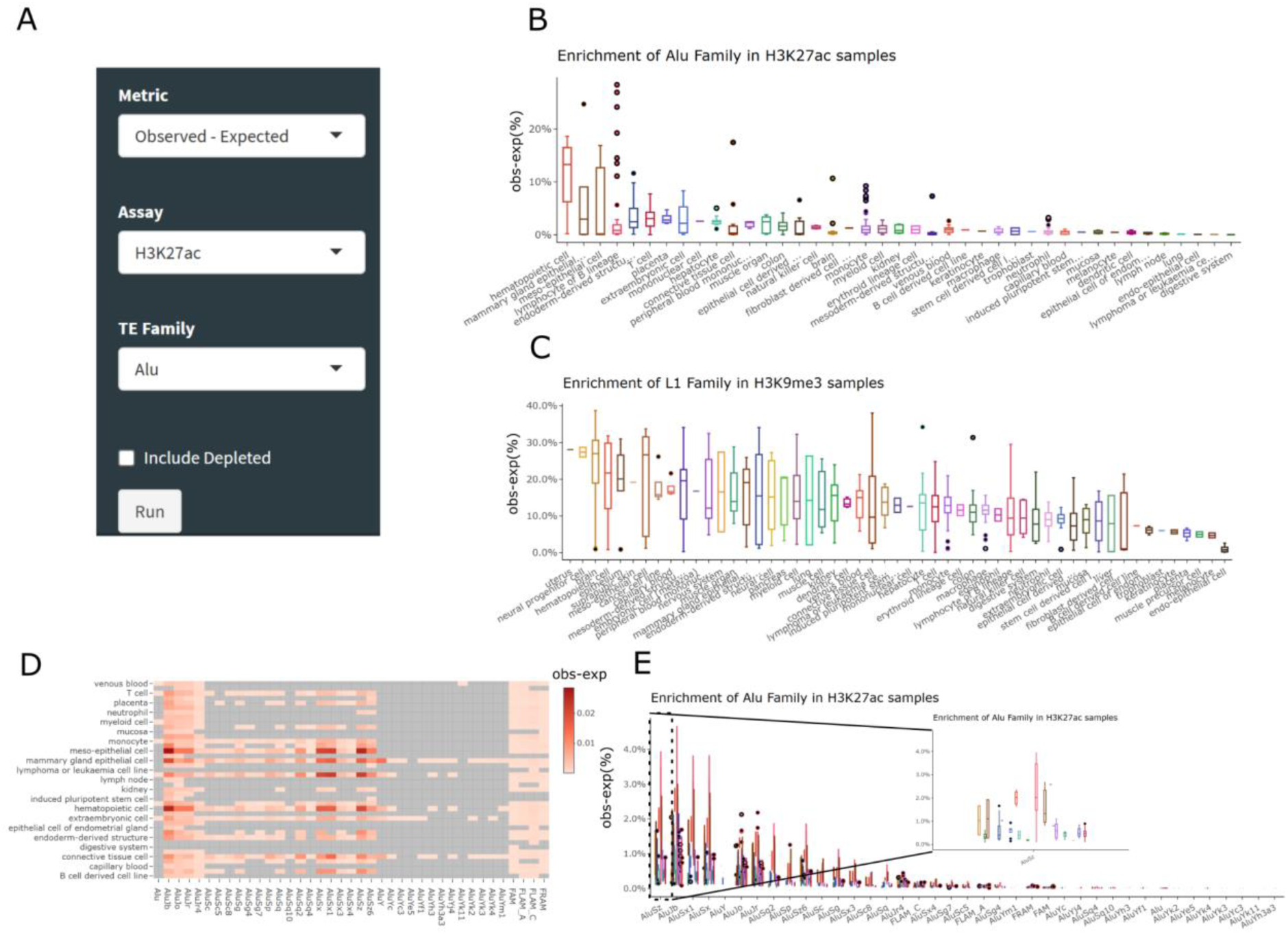
TE Subfamilies section shows the association between TE subfamilies and cell types. **A)** TE Subfamilies section input interface. The user must choose a metric, assay and TE family (here Alu). **B)** Boxplot of TE Family enrichment (obs-exp %) of selected TE family and histone (here Alu in H3K27ac) across cell types sorted in descending mean order. **C)** same as B for L1 in H3K9me3. **D)** TE subfamily enrichment heatmap of the TE subfamilies within a chosen family within cell types. **E)** TE subfamily enrichment boxplots within a chosen family (here Alu) across cell types (boxplot colors). Shown in zoom in is a specific subfamily (AluSz). The plots are interactive, and the user can zoom into select parts for more clarity.

Next, we report a heatmap of the TE enrichment (or count) for each subfamily within the selected TE family across the cell types. Through this plot, we can get a better idea of which TE subfamilies are enriched. For instance, with Alu and H3K27ac, we can discern that a few AluY subfamilies are not enriched, while a select few subfamilies such as AluSx and AluSz have higher enrichments (**Fig. 2D**). These results are also shown as boxplots to better show the distribution of enrichments across cell types (**Fig. 2E**). In this plot, we find that within AluSz specifically, the lymphocytes of B cells are the most enriched cell type but also have high variance. Finally, the data from the query can be explored or downloaded as a table (**Supplemental Fig 2C**).

Through this section of TEExplorer, users can gain insight on the level of enrichment of a chosen TE across cell types and view a breakdown of the TE subfamilies contributing to this enrichment.

### Importing user data for additional TE analyses and comparisons

The Import section allows users to upload their own samples and get their TE overlap as well as TE enrichments and comparisons to the EpiATLAS data. The user can upload multiple BED files and select which cell type and assay to compare them to (this would usually be from the same cell type as their samples, **Fig. 3A**). To better compare to the EpiATLAS data, an option to resize the uploaded sample’s peaks to 200bp (as was done to the EpiATLAS samples) is available. To demonstrate the features of this section of TEExplorer, we uploaded a T cell H3K9me3 sample that is already part of the EpiATLAS dataset. When run, TEExplorer measured the uploaded file’s peak overlap with TE annotations from RepeatMasker and compared it to EpiATLAS pre-generated random controls (matching the cell type and assay query). TEExplorer also projected the sample onto a UMAP plot to see if it aligned with the EpiATLAS data as a visual quality control (**Fig. 3B**). As expected, we can see that our sample clustered with both the H3K9me3 samples and the T cell samples and was very close to its duplicate sample within the EpiATLAS dataset (**Supplemental Fig 3A**).

**Figure 3.**
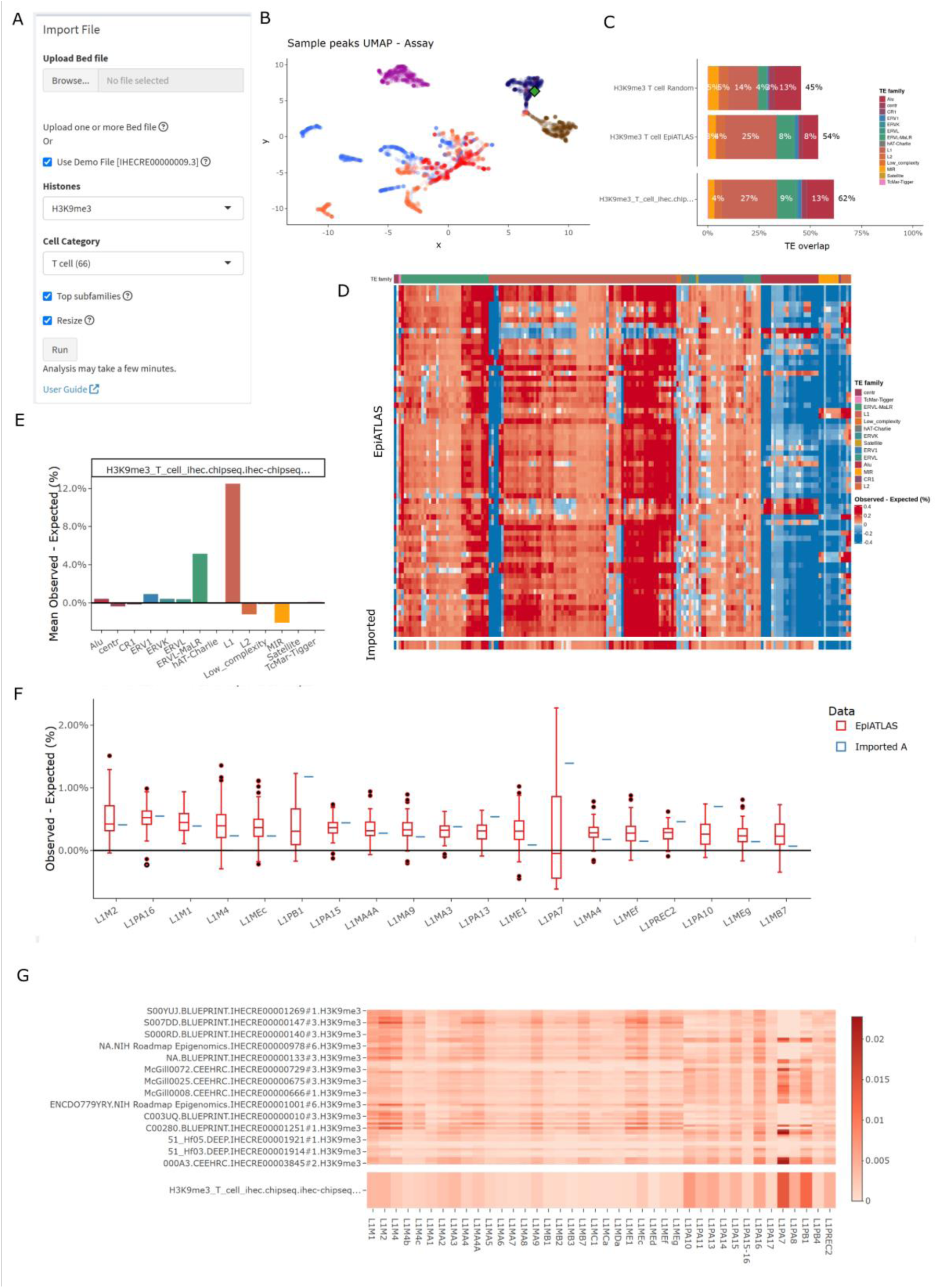
Import section provides an overview of TE enrichments within user data. **A)** Interface of data upload and input selection. **B)** UMAP of EpiATLAS samples based on peak counts within select 10kbp windows across the genome. Colored by assay. Green diamond is the uploaded sample’s overlayed predicted position. **C)** Bar plot of TE overlap the EpiATLAS data from selected histone and cell type and associated random simulation (top) and of the uploaded sample (T cell, H3K9me3, bottom) **D)** TE enrichment relative to random controls of select subfamilies from EpiATLAS data (top) TE enrichment relative to the mean of random controls of the uploaded samples. **E)** Mean enrichment relative to expected of TE families for each sample. **F)** Boxplot of the TE enrichment of TE subfamilies within a chosen family (here **L1**); comparison is between the EpiATLAS data (data) and the imported samples. **G)** heatmap of the TE enrichment of the TE subfamilies within a chosen family (here **L1**) within samples, shown in EpiATLAS samples (top) and the uploaded samples (bottom). The values shown in C, D and E can be Observed – Expected (shown), Observed count or Fold change. For all plots the EpiATLAS samples used are a subset matching the chosen cell type and assay.

This section of the tool also reports the TE overlap percentage by TE family for each sample as well as those from the mean of all EpiATLAS samples matching the selected cell type and assay. Our uploaded sample had much higher TE overlap (62%) than the mean from EpiATLAS T cells (54%) and the random controls (45%) (**Fig. 3C**). This higher TE overlap seemed to mostly come from Alu elements. The user can then choose between 3 metrics for the other results plots: “*Observed – Expected*”, “*Observed count*” or “*Fold change*”. The web tool then displays a heatmap of the TE subfamilies for the uploaded samples in comparison to the EpiATLAS samples matching the query (**Fig. 3D**).

There is a highlight subsection that displays the most extreme elements (enriched or depleted) to give an idea of the TEs that might be notable within the uploaded samples. It also shows the TEs with the largest difference from the mean of the selected EpiATLAS samples, which is another way to identify TE subfamilies that may be noteworthy within the imported samples. Then, we show a plot of the mean TE family enrichment or mean observed count within each uploaded samples (**Fig. 3E**), and then two TE subfamilies breakdown as boxplots (**Fig. 3F**) and a heatmap (**Fig. 3G**), once again comparing the user data to the EpiATLAS samples. Finally, we report a table of the data which includes the TE count for each subfamily per sample, the total peak count within the sample, the expected count according to the chosen cell type and assay, and two enrichment metrics. This table can be downloaded to power further analysis or allow custom plotting.

This section of TEExplorer allows users to get an overview of the TE content of their own data and compare it to the existing EpiATLAS data. It can give an idea of which TE families and subfamilies are most present, and if they are found more than expected. If all cell types are chosen, it can also be used to identify which cell type the uploaded cell type is most similar to.

### TEExplorer detects TE enrichment differences within flu infected macrophages

For a practical use case of TEExplorer, we investigated an independent dataset of monocyte-derived macrophages (MDMs) infected or not by Influenza A virus (IAV, commonly known as flu)^21^. We aimed to compare those samples with those of EpiATLAS and see if we could detect any difference between the infected and not infected samples. We uploaded a set of 8 MDM H3K27ac samples from the flu dataset: 4 infected (FLU) samples and 4 not infected (NI) samples (**Fig. 4A**). We used H3K27ac as our comparison assay, “macrophage” as our cell type, and enabled the option which resized peaks (**Supplemental Fig 4A**). After having run the tool, we assigned the FLU samples to group A and NI samples to group B (**Supplemental Fig. 4B**). Based on our quality control UMAP plot, we found that the uploaded samples did cluster with the EpiATLAS H3K27ac samples and the macrophage samples (**Fig. 4B, Supplemental Fig. 4C**), as we would expect.

**Figure 4.**
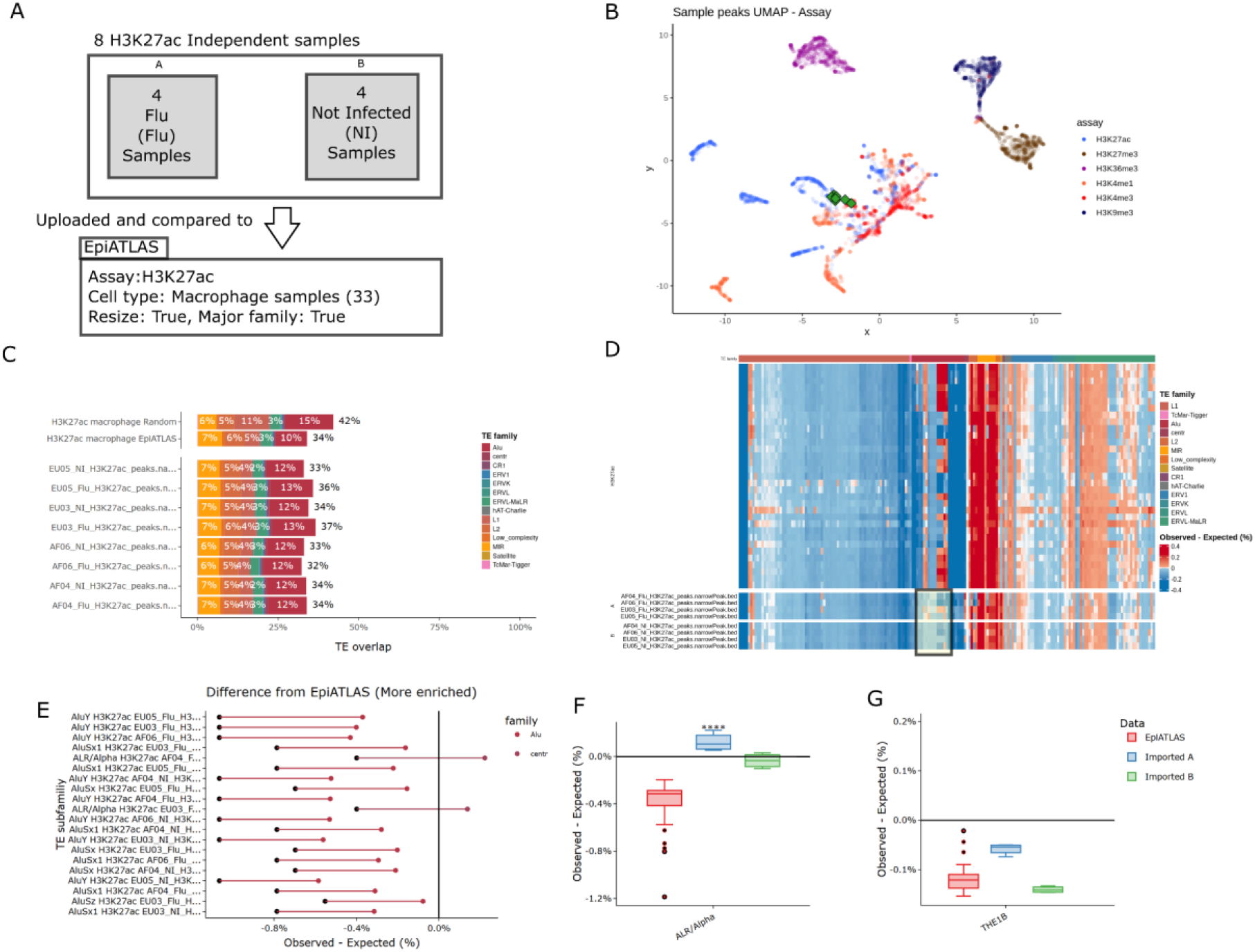
Analysis of TEs within Flu samples. **A)** Diagram of the data used and the portal’s input values (assay: h3k27ac, cell type macrophage, resize true) **B)** UMAP of EpiATLAS samples based on peak counts within select 10kbp windows across the genome. Colored by assay. The green diamonds are the uploaded sample’s overlaid predicted positions. **C)** Bar plot of TE overlap with the EpiATLAS data from selected histone and cell type and associated random simulation (top) and of the uploaded samples (bottom) **D)** TE enrichment relative to random controls of select subfamilies from EpiATLAS data (top) TE enrichment relative to the mean of random controls of the uploaded samples. In the highlighted rectangle are Alu elements distinct from EpiATLAS. **E)** Largest difference between uploaded samples and EpiATLAS samples. Black point is the mean EpiATLAS enrichment, colored point is the uploaded sample enrichment, bar in between represents the difference. **F)** Close up on boxplot of the TE enrichment (obs-exp %) of ALR/Alpha repeat (from centr family) compared between the EpiATLAS data and the uploaded samples (imported; A Flu, B non-infected). Stars are the significance of Wilcoxon test comparison **** = pval< 1e-04 **G)** same as F for ERVL-MaLR’s THE1B subfamily.

We found that our uploaded samples had between 32% to 37% TE overlap, which is in line with the 34% mean of the EpiATLAS H3K27ac macrophage samples and a bit lower than 42% for an adjusted random control (methods, **Fig. 4C**). There wasn’t any noticeable difference between the NI and FLU samples in terms of TE overlap. The TE family representation was also similar between the uploaded samples and EpiATLAS. When we looked in terms of enrichment relative to random control, we generally found the same patterns of enrichments across TE subfamilies, except a few Alu subfamilies that were noticeably less enriched than in EpiATLAS samples (**Fig. 4D**). There were more disparities between the uploaded samples and EpiATLAS when using the fold change metric (**Supplemental Fig. 5**).

The highlight sub-section shows that MIR, G-Rich and L2 TEs and repeats have the highest enrichments (**Supplemental Fig. 6A**), whereas the most depleted subfamilies were AluY and some L1PA elements (**Supplemental Fig. 6B**). The next plot highlights the TE subfamilies for which there are the most extreme differences between the uploaded samples and EpiATLAS (**Fig. 4E**). They were mostly Alu with ALR/Alpha elements being slightly enriched but depleted within EpiATLAS macrophages.

Within the Alu family, we identified AluY, AluSx, and AluSz as subfamilies that were more enriched than in EpiAtlas data, although they were still slightly depleted overall (**Supplemental Fig. 6C**). We also found that ALR/Alpha was more enriched in the uploaded samples and within those, more enriched for FLU samples than NI (**Fig 4F**). Within the ERVL-MaLR, the various THE1 subfamilies had a noticeable difference in enrichment between FLU and NI samples discernable through banding within the heatmap (**Supplemental Fig. 6D)**. Through the boxplots, the higher THE1B enrichment within FLU samples can be better observed (**Fig 4G**). A similar enrichment of THE1B within these influenza-infected samples was also observed by Chen et al.^10^, which shows that TEExplorer can recover results obtained independently.

Thus, TEExplorer could seamlessly display the TE overlap of our ChIP-seq samples, their TE enrichments, and comparisons with existing comparable samples. The results showed that the independent samples were like those within the EpiATLAS dataset, and that some significant differences could be observed between flu and non-infected samples.

## Discussion

TEExplorer provides an atlas of TE associations with histone marks and an easy-to-use interface for the exploration and interpretation of these results. In a previous study^4^, we found more associations between TE, histone mark, and cell type than we could highlight. TEExplorer makes these results available in an accessible and interpretable way, enabling researchers with an interest in a particular histone mark, cell type or TE to explore a vast array of associations directly.

Notably, the tool can also provide researchers with the TE overlap and enrichment of their own ChIP-seq BED files and show how it compares to the portal’s data and the simulated controls used for the portal’s analysis. We demonstrated the TE analysis abilities of our tool using independent H3K27ac flu and non-infected macrophage samples. We observed similarities between the TE enrichment profile of the uploaded macrophage samples and those from EpiATLAS but some notable differences could also be detected. The tool found that the most enriched TEs were from the MIR TE family, and the notable Alu enrichment was mostly from the AluY, AluSx and AluSz subfamilies. This was all done from an internet browser, without any installed tool or TE-specific knowledge.

However, there remain some limitations to our tool’s results. First, TE overlaps of the EpiATLAS dataset were done on data that did not keep multi-map reads. Thus, we may be losing TE instances from multi-map reads and under-estimating our TE overlap measurements for the younger families. Polymorphic and complex TEs that are not in the reference would also be missed. However, we do report a mappability estimate so that users can know if mappability could be impacting the results. In addition, since the EpiATLAS data comes from a group of consortiums, there may be heterogeneity across samples of the same cell type and technical artifacts. Finally, in order to minimize live calculations and runtime, the uploaded samples enrichments are calculated relative to expected observations from pre-generated EpiATLAS associated random simulation data. Thus, the enrichment analyses would perform best when the uploaded data is similar to the EpiATLAS data.

Despite these limitations, the portal provides more than 6.5 million TE overlap measurements and their associated enrichments for 4614 samples covering 57 cell types and 6 histone marks. The data can be queried according to TE, cell type and/or assay, is presented by multiple dynamic plots, and can be downloaded.

In summary, TEExplorer simplifies TE consideration by allowing researchers without TE expertise to explore which TEs are predominant within their experiment and how these TEs compare to expected levels. Thus, the portal can highlight TEs that may have been overlooked as causal elements within an experiment. The portal also performs simple analysis and reports TE overlap and TE enrichments within user uploaded data. While specialized tools will always be necessary for more in-depth TE analysis, our portal establishes a straightforward starting point towards greater TE consideration in epigenomic research. TEExplorer is currently available at: https://teexplorer.c3g.sd4h.ca

## Methods

### Implementation and data transformation

The webtool is powered by R^22^ and Shiny^23^ and is available online at https://teexplorer.c3g.sd4h.ca. To improve performance, tabular data were converted into SQLite databases^24^.

For quality control purposes and to compare samples, we kept a sample peak approximation in the form of the total peak count within 10 kilobase pair (kbp) windows across the genome. These counts were tabulated into a table of 321186 rows per sample which was reduced to the 20 000 most variable windows by variance (excluding the top 1000) and the index of these selected windows were saved to reuse on user uploaded data. We ran a UMAP^25^ (using the *umap* R package) dimension reduction on those 20000 windows for all samples and used the first 2 dimensions in a plot that shows sample similarity. We retain the resulting model for reuse with user uploaded data projection.

### Uploaded data source

The test sample of T cell H3K9me3 was taken from the IHEC EpiATLAS integrative analysis SFTP server on January 23rd 2023 and is also available on the IHEC data portal (https://epigenomesportal.ca/ihec/, https://ihec-epigenomes.org/epiatlas/data/, EPIRR: IHECRE00000009.3). The flu samples were obtained from the data of Aracena et al.^21^. The samples were converted from hg19 to hg38 using *liftOver*. A subset of 8 samples was used for our experiment (AF04_Flu_H3K27ac, AF04_NI_H3K27ac, AF06_Flu_H3K27ac, AF06_NI_H3K27ac, EU03_Flu_H3K27ac, EU03_NI_H3K27ac, EU05_Flu_H3K27ac, EU05_NI_H3K27ac).

### EpiATLAS data TE overlap, random control and enrichment

Histone ChIP-Seq peaks from the EpiATLAS dataset (downloaded from the SFTP server on January 23rd 2023 and also available on the IHEC data portal [https://epigenomesportal.ca/ihec/, https://ihec-epigenomes.org/epiatlas/data/]) were annotated using the UCSC RepeatMasker track^20^. We resized the peaks to 200bp around their center and counted the number of times the peaks overlapped TE instances with *bedtools*^26^ *intersect*. When more than one TE overlapped a peak, we selected the largest overlap. To have a random baseline to compare against, we generated, for each sample, a simulated library of 200bp random region with the same distribution of distance to nearest gene TSS. The distribution came from the following categories: TSS (within 1000 bp from TSS), Promoter (within 5000 bp upstream of TSS), Intragenic (Overlapping genes), Proximal (within 10 kbp from TSS), Distal (within 100 kbp from TSS) and Desert (further than 100 kbp from TSS). For each sample, the simulation was repeated 1000 times and we counted the number of times the observed count (from the sample) was higher than random baseline (“expected count”, current simulation) for each TE subfamily. A TE subfamily was identified as significantly over-represented (enriched) when it was higher in observed than random more than 995 times out of 1000 (p<0.005). Overlap percentages were measured as *peaks overlapping TE/sample peak count*. The “Observed – Expected” metric was calculated by subtracting the expected count (the mean overlap count across the 1000 simulation) from the observed overlap of the sample. Significant Observed – Expected (with depleted TEs excluded) was calculated using only the cases where the TE subfamilies were significantly enriched (and thus is always positive).

### Comparison between cell types

We took the table of TE enrichment measurements (excluding Simple_repeat family) and selected entries only fitting the desired context (input TE families and histone marks). If include depleted was not selected, we only used the TEs that were significantly enriched from the simulations as described in *EpiATLAS data TE overlap, random control and enrichment* section. From this point we had the TE enrichments of each TE subfamilies of given families and histone mark samples across all cell types. For TE family level measurements, we grouped by sample, cell type and TE family and summed the observed-expected measurements such to obtain one total TE family measurement per sample (different cell types).

### Estimation of TE mappability

TE mappability was estimated using the 50 bp Mappability track from the UCSC Genome browser, which is a conservative estimate because most of the reads of EpiATLAS had were longer and mappability increases with length. The coverage of all TE instances by the 50bp Unique Mappability (Umap 50) track gave us the proportion of each TE that could be uniquely mapped, which we used as a mappability metric.

### Data import transformation and analysis

TEExplorer can analyze BED and BED-like files up to 50 MB, e.g., narrow or broad peaks. The human reference used is hg38. For each uploaded hg38 BED file, an overlap with the RepeatMasker TEs and with a template BED file of 10kbp windows across the genome are performed using *genomicranges*^27^ and counts of peaks overlapping TEs and 10kbp windows respectively are obtained. Of the 321186 10kbp windows, the 20 000 most variable windows (excluding the top 1000) from our data, as previously determined and saved, are kept. To compare the user uploaded samples to those from our data, we project the user samples (with the R *predict* function applied to the retained UMAP model) onto the previously generated calculated UMAP plot on the EpiATLAS data.

The TE overlapping counts of uploaded samples are kept as a separate table with proportions being calculated through *peak overlapping TE count/peak count in sample* for each subfamily.

For the TE enrichments and comparisons between user and EpiATLAS data, a new temporary table is created by combining a subset of the EpiATLAS dataset that matches the chosen cell type and assay. The subset’s expected counts are averaged across all samples and used as the user uploaded sample expected TE counts. This provides a random sample approximation without having to perform the computation of all simulations. For the boxplots, a Wilcoxon test is performed between all groups with R function *stat_compare_means(label = “p.signif”, hide.ns = TRUE)* and default parameters.

### Data visualization

The plots were generated with *ggplot2*, *plotly* for dynamic plots, and *complexheatmap*^28^ for the static heatmaps. The interface was powered by *shinydashboard* and *shinyWidgets*^29^.

## Acknowledgements

This work was supported by a Canadian Institutes of Health Research (CIHR) program grant (CEE-151618) for the McGill Epigenomics Mapping Center, which is part of the Canadian Epigenetics, Environment and Health Research Consortium (CEEHRC) Network. G.B. is supported by a Canada Research Chair Tier 1 award and a FRQ-S, Distinguished Research Scholar award. We would like to acknowledge Calcul Quebec, SecureData4Health and the Digital Research Alliance of Canada for access to computing resources. We thank the IHEC consortium for their EpiATLAS dataset.

## Supplemental Figures

**Supplemental Fig 1:**
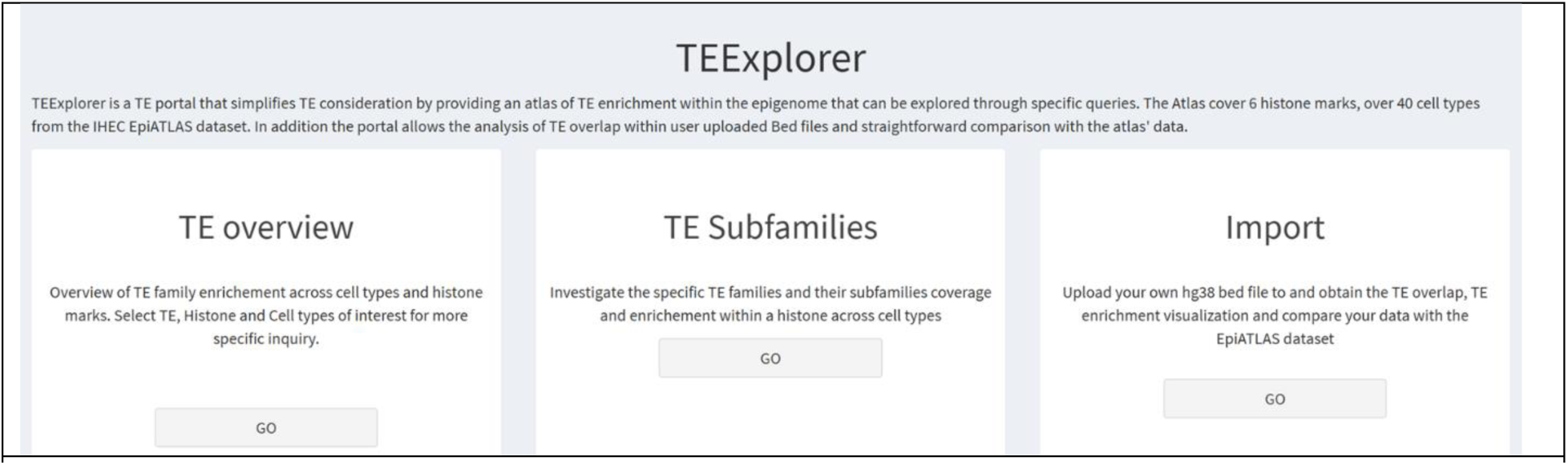
Web tool homepage and features. Screenshot of the home page showing TEExplorer’s 3 main sections.

**Supplemental Fig 2:**
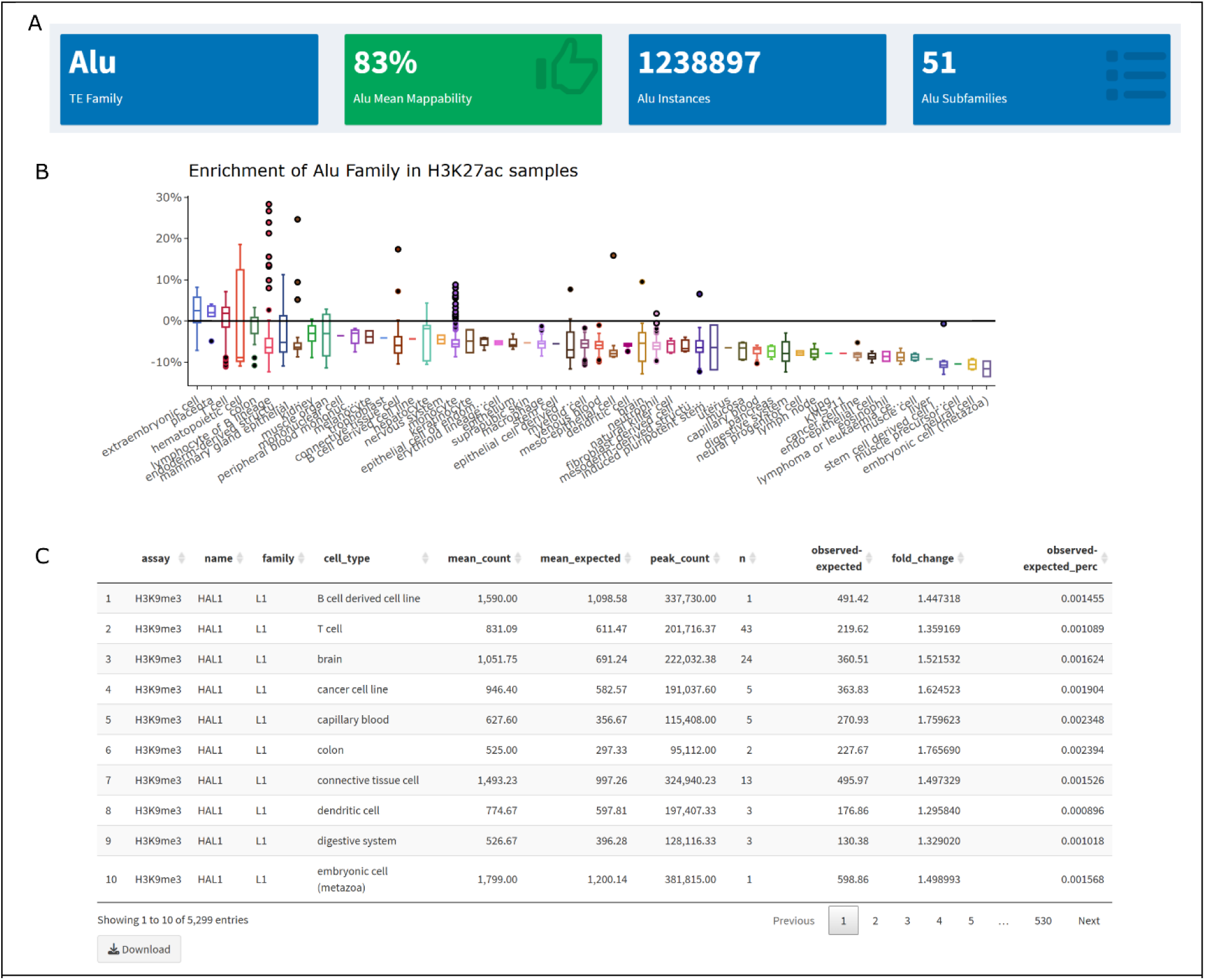
Subfamily section additional plots and information. **A)** Information on the selected TE family. The TE family, an estimate of its mapability, the number of Family enrichment (obs-exp %) of selected TE family and histone (here Alu in H3K27ac) Including non-enriched TEs across cell types sorted in descending mean order. **C)** Table of enrichments and counts for the TE and assay query across cell types.

**Supplemental Fig 3:**
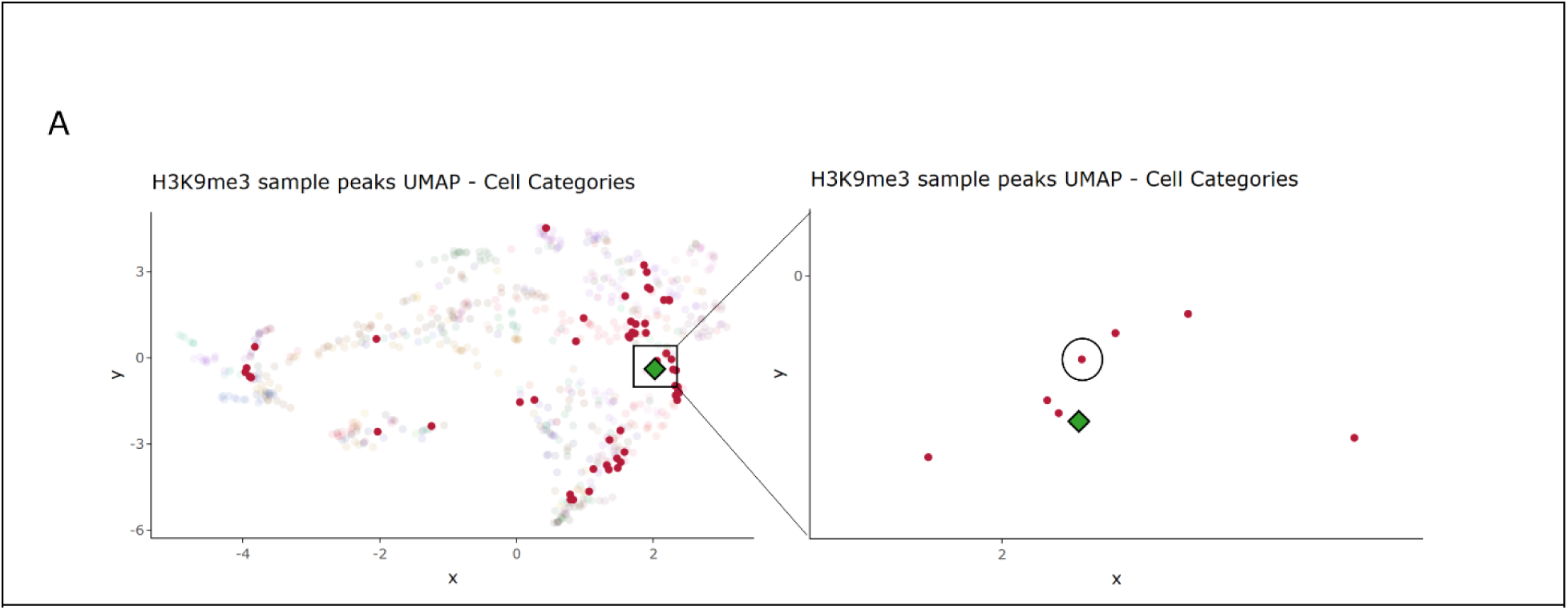
Import metric selection. **A)** UMAP of EpiATLAS samples based on peak counts within select 10kbp windows across the genome. Only H3K9me3 samples used. Colored by cell type. Green diamonds are the uploaded samples’ overlaid predicted positions. Opaque red points are the EpiATLAS T cell samples. Zoom upon a small selection around imported sample. Circled sample is the same sample as the uploaded (IHECRE00000009.3)

**Supplemental Fig 4:**
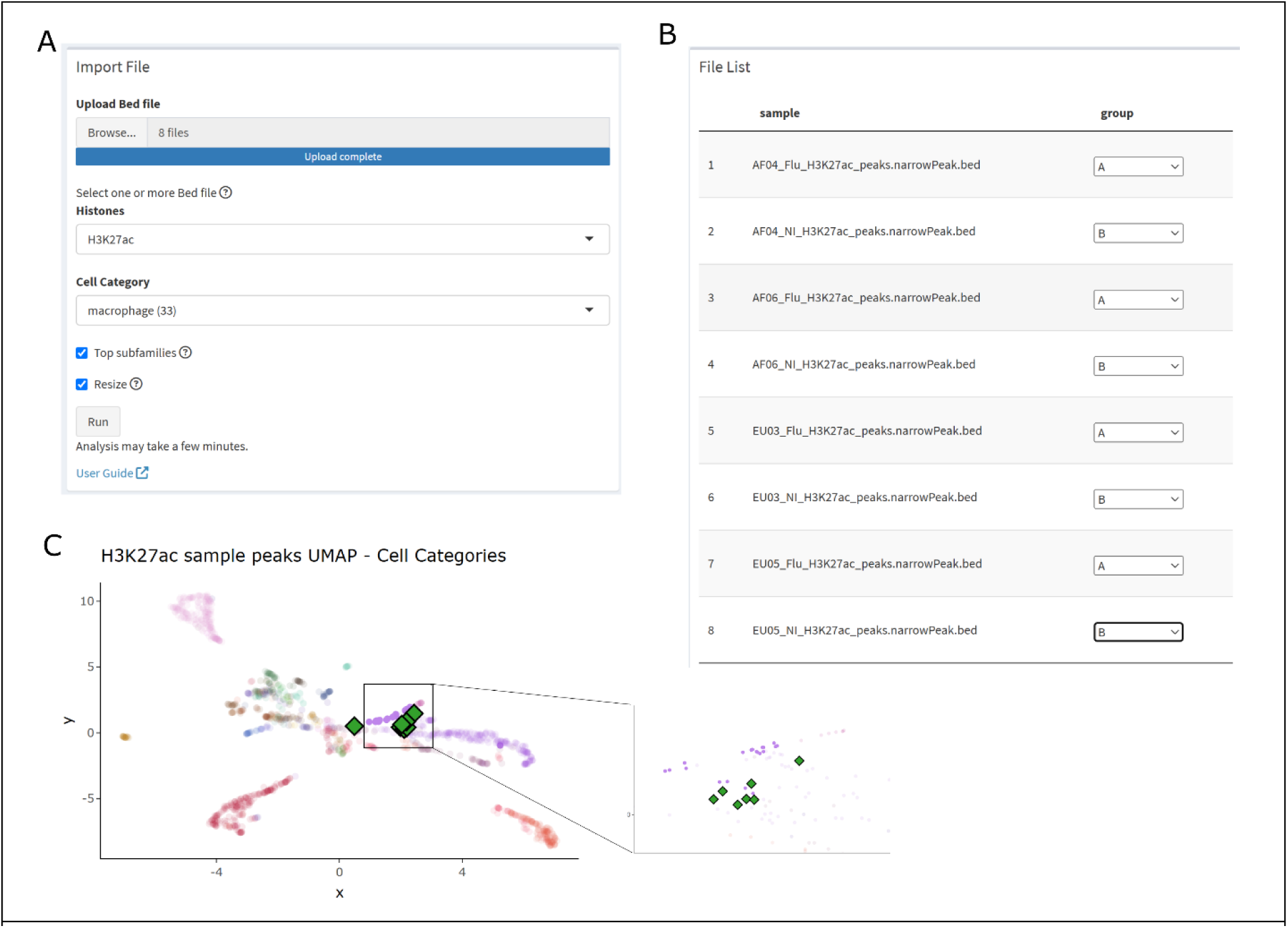
Flu data import input and additional QCA) Interface of data upload and input selection. **B)** File list grouping. Assignment of samples to groups. Flu samples assigned A, not infected assigned B. **C)** UMAP of EpiATLAS samples based on peak counts within select 10kbp windows across the genome. Only H3K27ac samples used. Colored by cell type. Green diamonds are the uploaded samples’ overlaid predicted positions. Opaque Purple points are the EpiATLAS macrophage samples. Zoom upon a small selection where the imported samples were concentrated.

**Supplemental Fig 5:**
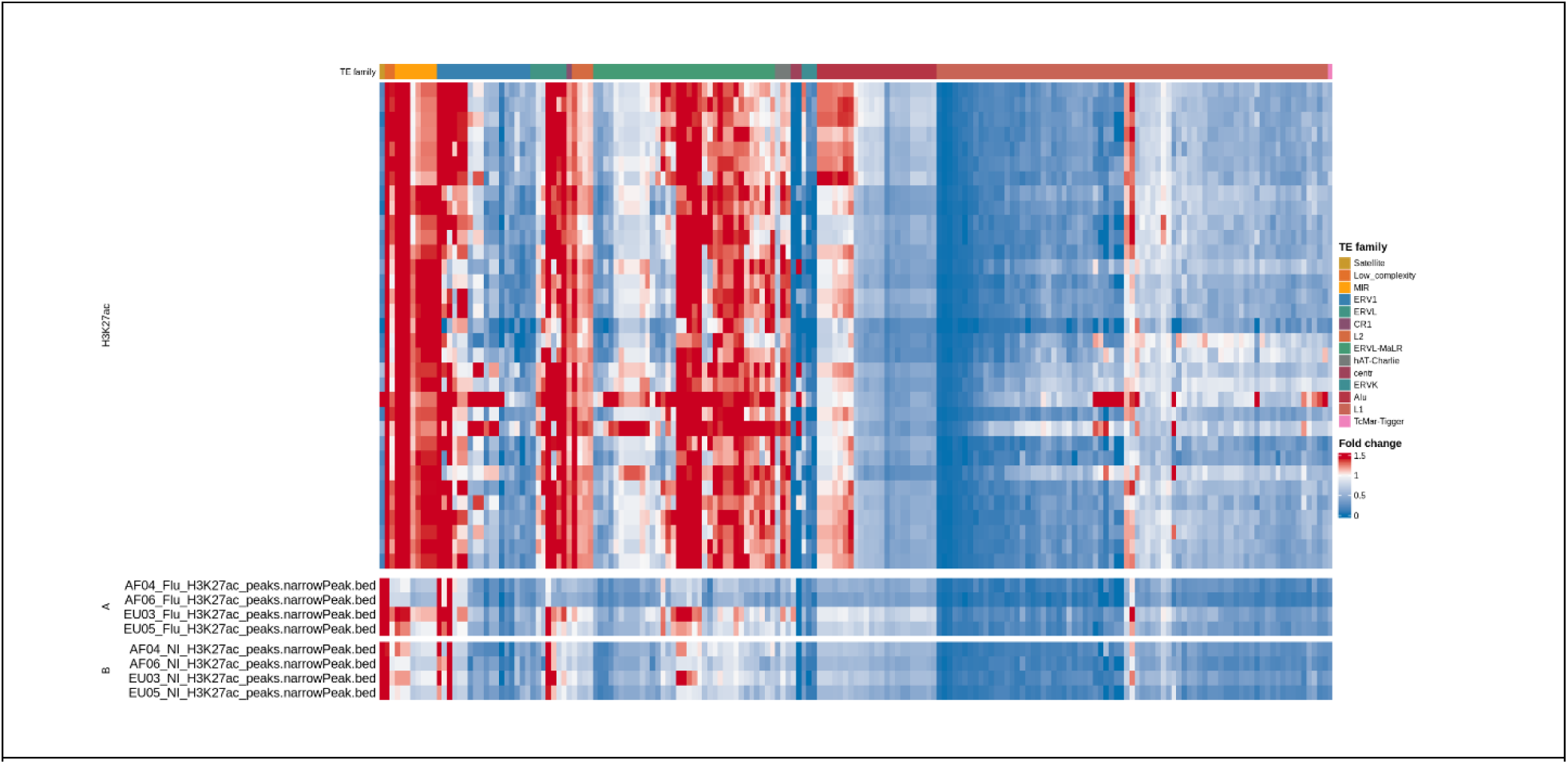
Flu data Import alternate metrics heatmaps. TE enrichment (fold change) relative to random controls of select subfamilies from EpiATLAS data (top) TE enrichment (fold change) relative to the mean of random controls of the uploaded samples grouped by letter group (bottom).

**Supplemental Fig 6:**
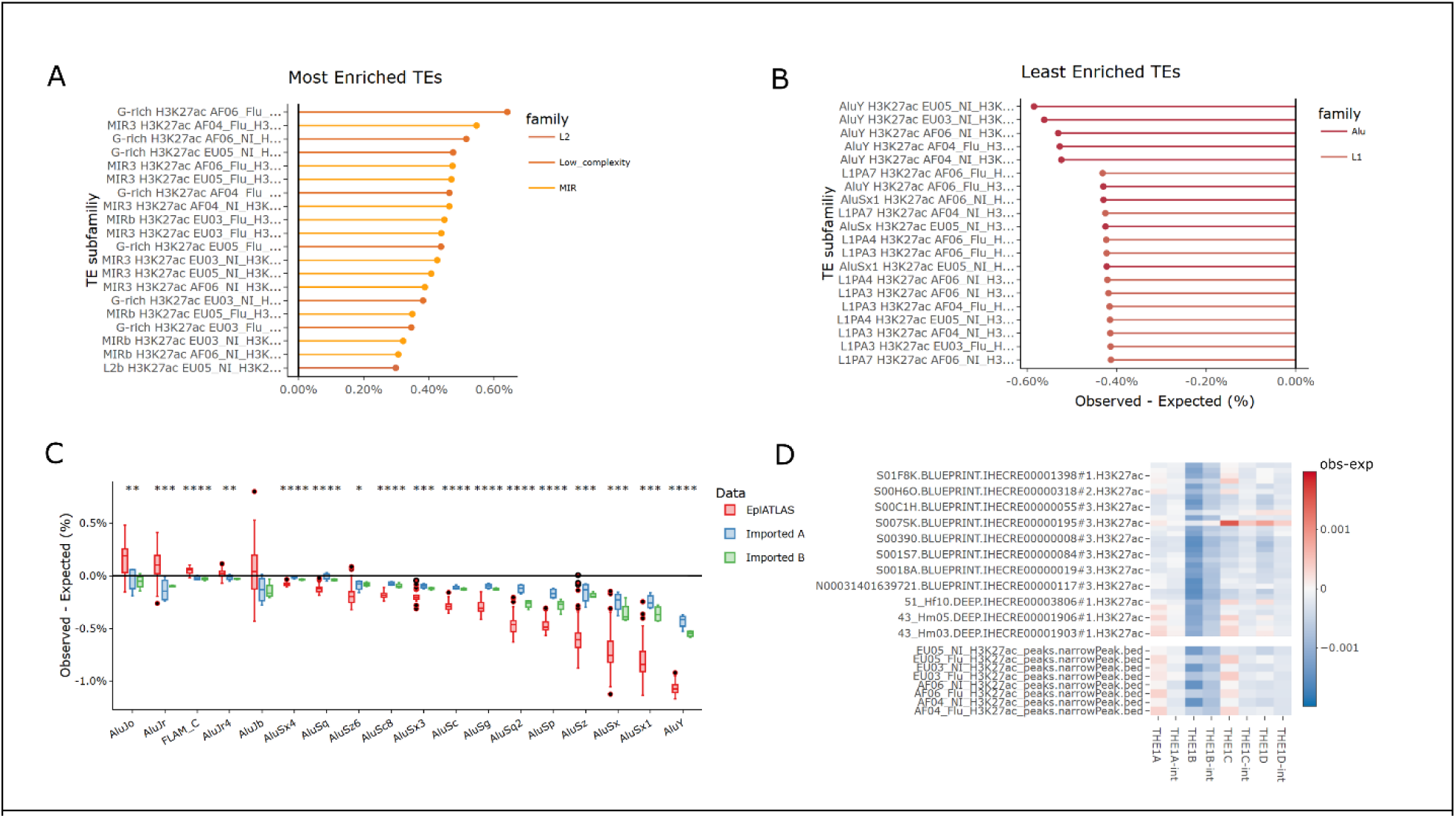
Flu data import highlights. **A)** Top 20 most enriched TE subfamilies within the 8 imported samples. according to observed-exp (%) colored by TE family. **B)** Same as A for the least enriched (most depleted) TEs. **C)** Boxplot of the TE enrichment (obs-exp) of TE subfamilies within a chosen family (here Alu) comparing the EpiATLAS data (data) and the uploaded samples (imported). Stars are the significance of Wilcoxon test comparison **** = pval< 1e-04; *** = pval<0.001; ** = pval<0.01; * = pval<0.05 **D)** heatmap of the TE enrichment (obs-exp) of the TE subfamilies within a chosen family (here ERVL-MaLR) within samples. Shown are EpiATLAS samples (top) and the uploaded samples (bottom).

## References

1. Wells, J. N. & Feschotte, C. A Field Guide to Eukaryotic Transposable Elements. Annu Rev Genet 54, 539–561 (2020).

2. Bourque, G. et al. Ten things you should know about transposable elements. Genome Biology 19, (2018).

3. Jacques, P.-É., Jeyakani, J. & Bourque, G. The Majority of Primate-Specific Regulatory Sequences Are Derived from Transposable Elements. PLoS Genet 9, e1003504 (2013).

4. Hyacinthe, J. & Bourque, G. Transposable elements impact the regulatory landscape through cell type specific epigenomic associations. 2024.08.07.606967 Preprint at 10.1101/2024.08.07.606967 (2024).

5. Pehrsson, E. C., Choudhary, M. N. K., Sundaram, V. & Wang, T. The epigenomic landscape of transposable elements across normal human development and anatomy. Nat Commun 10, 1–16 (2019).

6. Bourque, G. et al. Evolution of the mammalian transcription factor binding repertoire via transposable elements. Genome Res. 18, 1752–1762 (2008).

7. Su, M., Han, D., Boyd-Kirkup, J., Yu, X. & Han, J.-D. J. Evolution of Alu Elements toward Enhancers. Cell Reports 7, 376–385 (2014).

8. Jjingo, D. et al. Mammalian-wide interspersed repeat (MIR)-derived enhancers and the regulation of human gene expression. Mobile DNA 5, 14 (2014).

9. Bogdan, L., Barreiro, L. & Bourque, G. Transposable elements have contributed human regulatory regions that are activated upon bacterial infection. Philosophical Transactions of the Royal Society B: Biological Sciences 375, 20190332 (2020).

10. Chen, X. et al. Transposable elements are associated with the variable response to influenza infection. Cell Genomics 3, (2023).

11. Goerner-Potvin, P. & Bourque, G. Computational tools to unmask transposable elements. Nature Reviews Genetics 19, 688–704 (2018).

12. Wilkinson, M. D. et al. The FAIR Guiding Principles for scientific data management and stewardship. Sci Data 3, 160018 (2016).

13. McLean, C. Y. et al. GREAT improves functional interpretation of cis-regulatory regions. Nat Biotechnol 28, 495–501 (2010).

14. Kent, W. J. et al. The Human Genome Browser at UCSC. Genome Res. 12, 996–1006 (2002).

15. The ENCODE Project Consortium. An integrated encyclopedia of DNA elements in the human genome. Nature 489, 57–74 (2012).

16. Lonsdale, J. et al. The Genotype-Tissue Expression (GTEx) project. Nat Genet 45, 580–585 (2013).

17. Fernandes, J. D. et al. The UCSC repeat browser allows discovery and visualization of evolutionary conflict across repeat families. Mobile DNA 11, 13 (2020).

18. Shen, J. et al. Exploring the epigenome profiles of repetitive elements with the WashU Repeat Browser. Genome Res. gr.279764.124 (2025) doi:10.1101/gr.279764.124.

19. Bujold, D. et al. The International Human Epigenome Consortium Data Portal. cels 3, 496–499.e2 (2016).

20. Smit, A. F. A., Hubley, R. & Green, P. RepeatMasker Open-4.0. (2013).

21. Aracena, K. A. et al. Epigenetic variation impacts individual differences in the transcriptional response to influenza infection. Nat Genet 56, 408–419 (2024).

22. R Core Team. R: A language and environment for statistical computing. R Foundation for Statistical Computing,.

23. Winston Chang, et al. shiny: Web Application Framework for R. (2026).

24. Müller, K., Wickham, H., James, D. A. & Falcon, S. RSQLite: SQLite Interface for R. (2024).

25. McInnes, L., Healy, J. & Melville, J. UMAP: Uniform Manifold Approximation and Projection for Dimension Reduction. Preprint at http://arxiv.org/abs/1802.03426 (2020).

26. Quinlan, A. R. & Hall, I. M. BEDTools: a flexible suite of utilities for comparing genomic features. Bioinformatics 26, 841–842 (2010).

27. Lawrence, M. et al. Software for Computing and Annotating Genomic Ranges. PLOS Computational Biology 9, e1003118 (2013).

28. Gu, Z., Eils, R. & Schlesner, M. Complex heatmaps reveal patterns and correlations in multidimensional genomic data. Bioinformatics 32, 2847–2849 (2016).

29. Perrier, V., Meyer, F. & Granjon, D. shinyWidgets: Custom Inputs Widgets for Shiny. (2024).

